# Predicting Clinical Outcomes in *Helicobacter pylori-*positive Patients using Supervised Learning through the Integration of Demographic and Genomic Features

**DOI:** 10.1101/2025.07.28.667137

**Authors:** Venkatesh Narasimhan, Sreya Pulakkat Warrier, Jobin Jacob John, Monisha Priya T, Niriksha Varadaraj, Greeshma Grace Thomas, Balaji Veeraraghavan

## Abstract

**Background:** *Helicobacter pylori (H. pylori)* infection is widespread globally and is linked to outcomes ranging from chronic gastritis to gastric cancer. However, only a minority of infected individuals progress to malignancy, influenced by a mix of bacterial, host, and environmental factors. Current predictive approaches are limited due to relying mainly on clinical and lifestyle data. Genomic approaches have been sparsely used, and thus their incorporation into machine learning models could ensure early and personalized detection. This study aimed to evaluate the impact of integrating host metadata with genomic features from *H. pylori* to predict gastric cancer outcomes and identify associated variables.

**Methods:** 1,363 publicly available *H. pylori* genomes with associated host information between 1991 and 2024 were collected from NCBI and EnteroBase. Demographic features, virulence genes, sequence-derived and variant-based features were extracted. Machine learning models were then developed to classify infection outcomes into gastric cancer and non-gastric cancer. Logistic regression, an interpretable baseline model, was compared against higher-performance ensemble models (XGBoost, Random Forest). Model performance was assessed using recall, precision, AUROC, and AUPRC curves.

**Results:** The logistic regression model achieved a recall of 0.736 (95% CI: 0.644-0.831) for gastric cancer and an AUROC of 0.888 (95% CI: 0.843-0.929). Both XGBoost and Random Forest models outperformed the baseline model with AUROC values ranging from 0.950-0.954 (95% CI: 0.904-0.976). Black-box model recall for gastric cancer detection improved compared to the baseline by 8.3% for XGBoost (0.797, 95% CI: 0.711-0.877), and 11.4% for Random Forest (0.820, 95% CI: 0.734-0.896). Across models, patient age consistently emerged as the strongest predictor of gastric cancer, with several sequence-derived genomic features beyond pre-established virulence genes contributing to the infection outcome differences.

**Conclusion:** This study demonstrates that combining pathogen genomics with host demographics uncovers novel risk factors and ensures early detection with high predictive power. The use of explainability methods like SHAP allows for greater interpretability by clinical professionals and improves informed decision-making processes. Validation and translation into clinical practice can be carried out with broader, diverse datasets along with the inclusion of additional host and lifestyle variables.

## 1. Background

*Helicobacter pylori* (*H. pylori*) is a highly motile, helical and gram-negative bacterium that is known for colonising the human gastrointestinal tract, and primarily associated with chronic active gastritis. *H. pylori* infection is strongly linked to the development of serious gastrointestinal diseases and accounts for up to 90% of duodenal ulcers, and around 80% of stomach ulcers and 90% of gastric carcinoma and mucosa-associated lymphoid tissue (MALT) lymphoma cases^1^. In accordance with the burden posed by these diseases, the World Health Organisation (WHO) recommended prioritising *H. pylori* eradication strategies in 2014 to help in reducing stomach cancer mortality rates worldwide.

At the global level, *H. pylori* affects almost half of the world’s population, and has a disproportionately high prevalence of around 50.8% in resource-limited regions, which are often characterized by suboptimal sanitation and healthcare access ^2^. Gastric cancer is also relatively uncommon among younger individuals with fewer than 10% of cases occurring before the age of 45. There is a sharp spike in the lifetime risk of gastric cancer for infected individuals, with the probability being nearly double compared to uninfected individuals^3^.

The clinical outcomes of *H. pylori* infection considerably vary across geographic populations and are influenced by a variety of factors like bacterial genotype diversity, host immune responses, and environmental factors. The bacterium’s ability to thrive in the acidic gastric environment has been linked to specialized virulence mechanisms. These help it facilitate persistent colonization and chronic inflammation, which dictate the host’s gastritis phenotype and disease trajectory. There are several *H. pylori* genes that have been implicated in the development of severe gastroduodenal diseases. The cytotoxin-associated gene A (*cagA)* gene, encoding the CagA protein is found at the end of the cag pathogenicity island (cagPAI), and is associated with increased cancer risk and severe infections^4^. The marker is more discriminative in Western countries on account of only around 60% of strains carrying *cagA*, compared to over 90% of East Asian strains being *cagA*-positive^5^. The vacuolating cytotoxin A (*vacA)* gene encodes the vacA cytotoxin which helps in inducing vacuole formation, autophagy modulation and evading immune responses^6,7^. Strains harbouring the *vacA* s1/m1/*cagA*+ genotype show a 4.8-fold increased risk of progressing to precancerous lesions compared to *vacA* s2/m2/*cagA*-strains^8^. The *babA2* gene, encoding the blood group antigen binding adhesin (BabA) that binds to mucosal Lewis^b^ blood group antigens, helps in aiding colonisation and increasing bacterial load^9^. The triple positive genotype possessing *babA2, vacA* s1 and *cagA* genes offer stronger gastric cancer discrimination than *vacA* and *cagA* alone^10^. The outer inflammatory protein A (*oipA*) gene promotes epithelial attachment, IL-8-mediated inflammation and disruption of cell turnover, and is often found co-expressed with other virulence genes^11–16^. *homB* gene has been found to aid adhesion and IL-8 induction, while *sabA* promotes neutrophil activation and inflammation through binding to sialylated Lewis antigens (sLe^X^)^17–20^.

The *htrA* gene is closely linked to tumor suppression and metastasis, and the *iceA1* gene has also been significantly associated with gastric cancer^21–23^. The *hopQ* type I allele is linked with increased gastric cancer and gastric ulcer risk, especially in East Asia where it is commonly present with *cagA*, while type II *hopQ* is more frequent in Western strains^24^.

Diagnosing *H. pylori* infection is reliant on a combination of methods. These include invasive methods such as endoscopy and biopsy, and non-invasive approaches like urea breath tests, stool antigen tests, and serological assays. The diagnostic tool choice is guided by a lot of factors like patient history, clinical presentation and resource availability. Typical eradication therapy usually involves a potent acid suppressant with antibiotics and bismuth compounds in some cases. Curbing the alarming rise of antibiotic resistance among *H. pylori* strains relies on judicious antibiotic use and resistance surveillance methods^25^.

Understanding the genetic diversity of *H. pylori* and its associated phenotypic variability can be used to build accurate prediction models using machine learning approaches. This could help improve therapeutic approaches by stratifying patients based on infection outcomes^26^.

Despite there being some studies on predicting gastric cancer in *H. pylori* infected patients, they have primarily relied on clinical and lifestyle data and often included patients without confirmed *H. pylori* infection^27–29^. To our knowledge, there have been no studies that have integrated clinical host metadata, aggregated variant annotation features from VCF files, virulence gene presence and absence data, and sequence-derived features from *H. pylori* genomes in a single, unified machine learning framework. In this study, we focused solely on *H. pylori* infected patients and developed a machine learning pipeline to classify gastric cancer versus non-gastric cancer cases using a combination of the above mentioned features. A comprehensive summary of the study workflow is provided in Figure 1. We aim to provide researchers with novel genomic markers that can be further analyzed for causation in wider studies, along with the validation of established causative factors of gastric cancer like age. We hope this will improve stratification of infection responses and prevent delayed care for high-risk patients, along with increasing physician trust and augmenting decision-making.

**Figure 1:**
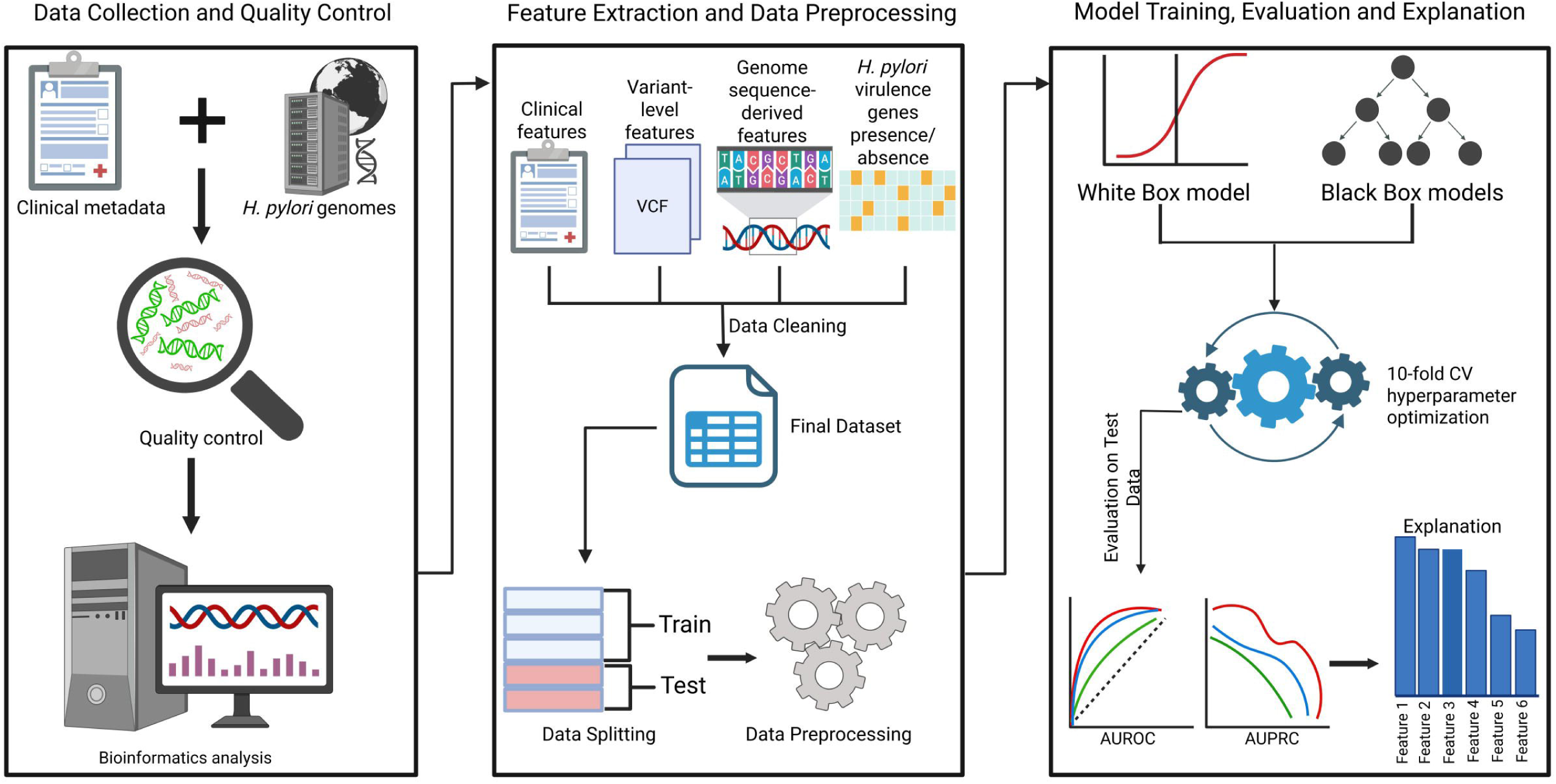
Overview of the machine learning workflow for classification of gastric cancer and non-gastric cancer using *H. pylori* genomes and clinical data. The workflow consists of three stages, depicted from left to right, with arrows indicating the flow of the stages: (1) Data collection and quality control, host clinical metadata and *H. pylori* whole genomes are integrated and filtered for quality, and bioinformatics analysis (variant calling); (2) Feature extraction and data preprocessing (normalization, one-hot encoding, label encoding), including clinical features, variant-level features from VCF files, and genome sequence-derived features and presence/absence data of *H. pylori* virulence genes to construct the final dataset, followed by data preprocessing; and (3) Model training, evaluation and explanation, involving binary classification using white-box (Logistic Regression) and black-box models (XGBoost and Random Forest) with 10-fold cross-validation and hyperparameter bayesian optimization, performance evaluation using AUROC and AUPRC metrics, and finally feature importance interpretation.

## 2. Methods

### 2.1 Data Collection

*Helicobacter pylori* genomes from NCBI and Enterobase were filtered for the presence of host-age, host-sex, host geographical location and disease phenotype information, resulting in 1587 genomes. The genomes from Enterobase were pre-assembled and case controls were excluded. NCBI Biosample Metadata was similarly downloaded and case controls and duplicates were excluded. Pre-assembled genomes from GenBank and RefSeq were downloaded using NCBI Datasets, while Sequence Read Archive (SRA) database sequences consisting of high-throughput short reads were downloaded through Fastq-dump from SRA Toolkit (version 3.2.1).

### 2.2 Genome Quality Control and Assembly

The unassembled short reads were assembled with the help of SKESA (version 2.4.0). Quality control was carried out using Pathogenwatch (version 23.2.1) to filter out contaminated genomes belonging to the wrong species, resulting in 781 Enterobase downloaded genomes and 582 genomes from NCBI. The final dataset consisted of 1,363 genomes which passed the quality control step and were collected in the years spanning from 1991-2024.

### 2.3 Feature Extraction

In order to develop a machine learning model well-equipped to predict gastric cancer status in *H. pylori* infected patients, a diverse set of features was included in the final dataset. These included clinical metadata from NCBI and Enterobase, gene presence and absence information obtained with the help of BLASTN-based homology searches, aggregated variant annotation features from VCF files and genome-derived sequence descriptors.

### 2.4 Clinical and Epidemiological features

The clinical metadata associated with each *H.pylori* genome was collected from Enterobase and NCBI databases (Supplementary Table S1). The age, sex, and geographical location of the host along with their disease phenotype was extracted. With the help of clinical expertise, this disease phenotype was converted into one of two labels: gastric cancer and non-gastric cancer. This served as the label for the machine learning model and was appropriately encoded as mentioned in later sections.

### 2.5 Gene Presence and Absence features

In addition to the clinical metadata, gene variant presence and absence information was also derived by identifying relevant gene variants associated with gastric and non-gastric cancer phenotype through exhaustive literature review.

These reference gene variant sequences linked to certain strains were extracted from a collection of genomes. These included cagA, babB, oipA, sabA and vacAs1m1 from *H.pylori* strain 60190; babA2 from *H.pylori* strain ATCC 43504; hopQI from *H.pylori* strain J99; homB, hrgA, htrA, iceA1 from *H.pylori* strain 26695; hopQII from Tx30a.

Each of these assembled genomes were formatted into a separate BLAST compatible database. Each gene sequence was queried against each of the 1363 genome databases with the help of BLASTN (version 2.12.0). A gene was considered present in a genome if the top hit satisfied the below criteria:

- Percentage identity >= 90%
- Query coverage >= 80%
- E-value <= 1e-5

The presence of each gene was encoded as 1 and the absence as 0, resulting in a binary feature set (Supplementary Table S2). These thresholds were selected in order to balance sensitivity and specificity and to ensure accurate homologous gene detection by minimizing false positives.

### 2.6 Assembled Genomic Sequence Derived features

To capture additional biological information, numerical sequence descriptors were extracted directly from the genomes using two Python-based packages, iFeatureOmega (version 1.0.2) and MathFeature (version 1.0) (Table 1).

**Table 1:** Feature types utilised in this study and the respective package.

| Package | Feature type |
| --- | --- |
| iFeatureOmega | Z-curve (144-bit) |
|  | Kmer Type 1 |
|  | RCKmer Type 1 |
|  | Nucleic Acid Composition (NAC) |
|  | Multivariate Mutual Information (MMI) |
|  | CKSNAP type 1 |
| MathFeature | Shannon Entropy |
|  | ORF-Based Features |

These sequence-based features were computed by the packages at a contig level, which were then averaged in a length-weighted manner to generate a single representative feature vector per genome (Supplementary Table S3).

Specifically, for a given feature *f*, the genome-level feature average *F* was calculated as:

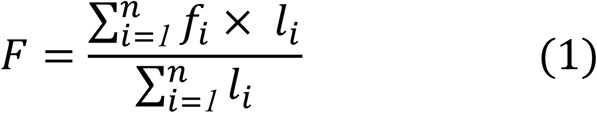

where:

- *f_i_* is the feature value for the *i*-the contig,
- *l_i_* is the length (in base pairs) of the *i*-th contig,
- and *n* is the total number of contigs for that genome.

This method ensured that longer contigs which typically provide more reliable sequence information would contribute proportionally to the final feature representation.

### 2.7 Variant-Based Features from VCF files

Variant level features were extracted from genome-wide mutation profiles to include impact about sequence variation and their functional effects. The strain 26695 was used as a reference to call variants using Snippy (version 4.6.0), a rapid bacterial variant calling pipeline, on the assembled *H.pylori* genomes. Following variant calling, the Snippy-created VCF files were passed to another tool known as bcftools (version 1.2.1) for identifying the number of SNPs and MNPs. Using bcftools’ (version 1.2.1) stats module, the raw counts of the following features were extracted SNPs, MNPs, Indels, and all possible nucleotide changes (e.g.,A → T, T → C, etc.). The variant effect functional annotations were also extracted from the VCF files to classify sequence changes based on their predicted effects on coding regions. The total counts of each variant effect were calculated per genome. These included types of mutations such as missense variants, frameshift mutations, mutations that affect start codons and combinations of these. For each genome, the number of variants falling into each functional category was counted and recorded as individual features.

These metrics were aggregated into per-genome features to help capture unexplored signals associated with the gastric cancer phenotype (Supplementary Table S4).

### 2.8 Data Preprocessing

A comprehensive series of steps were carried out to preprocess the dataset to make it amenable for model training (Supplementary Table S5). The versions of scikit-learn, Pandas, matplotlib and Python used were 1.6.1, 2.2.3, 3.10.1 and 3.12.3.

Minor corrections were carried out to standardise categorical entries including correcting typographical errors, standardising the case of the characters and removing entries where the host sex was indeterminate, resulting in 1362 genomes.

The dataset was then split into training sets and test sets using the StratifiedShuffleSplit module from scikit-learn (version 1.6.1) (80% training set, 20% test set). For the input features, preprocessing for the input features involved normalisation using the MinMaxScaler module and one-hot encoding using the OneHotEncoder module in scikit-learn.

Label encoding was used to convert the categorical class labels of ‘Gastric cancer’ and ‘Non-gastric cancer’ into numeric representations using the LabelEncoder module in scikit-learn. Since the LabelEncoder module encodes the classes in alphabetical order, ‘Gastric cancer’ was encoded as 0 and ‘Non-gastric cancer’ was encoded as 1.

The same processes of normalisation and encoding were also applied separately to the test set to ensure that the preprocessing steps were consistent across training and test datasets. The transformations performed on the test set were done independently to prevent any form of information leakage. The resulting dataset was rendered suitable for further feature selection and model training steps.

### 2.9 Feature Selection

Feature selection was only performed for the black-box models (XGBoost and Random Forest) to mitigate overfitting, enhance generalization and improve the model’s explainability. In contrast, the baseline model (Logistic Regression) was trained on the complete feature set to provide a consistent benchmark for comparison. The ShapRFECV module in the probatus package (version 3.1.2) which performed recursive feature elimination based on SHAP importance scores was used to select features. The features with the lowest SHAP scores were iteratively removed to get the final feature set.

The optimal feature subset was selected by maximizing the base model’s (LightGBMClassifier) performance used for performing feature selection. This resulted in a final set of 93 features, which was used to subset the original dataset.

### 2.10 Model Training and Development

A Logistic Regression classifier was trained as the baseline model and chosen for its white-box functionality and for its ease of interpretability. A random state of 26 was used to maintain reproducibility.

Class imbalance in the dataset was handled using SMOTE-NC. It was preferred over SMOTE due to its ability to handle both categorical and continuous features by generating synthetic samples for the minority class while retaining the categorical nature of categorical variables. It was applied within a generated pipeline to the training folds within each cross-validation split to ensure that synthetic samples did not influence the model evaluation step on the validation step. The full model training pipeline was built using imblearn’s make_pipeline function and had SMOTE-NC and Logistic Regression classifier (with random state=26 to maintain reproducibility) as its two components.

Cross-validation was carried out using StratifiedShuffleSplit by splitting the dataset into 10 folds for robust evaluation, with the stratification step preserving the class distribution during both model training and evaluation. Recall was selected as the scoring metric to prioritise accurate identification of gastric cancer cases and minimise any false negatives. After cross-validation, the model was refitted on the entire training dataset to optimise its performance before applying it to the test set.

An XGBoost classifier was trained as the black-box model and chosen for its black-box high performance. The steps taken for preprocessing the dataset were identical to the baseline model, including the chosen random state.

Categorical feature indices were identified and used with SMOTE-NC and a pipeline to first oversample and then train an XGBClassifier on the reduced feature set. Hyperparameter tuning was carried out using the BayesSearchCV function from scikit-optimize package, which employs Bayesian optimization to automatically search for the best hyperparameters to enhance the performance of the model.

Probability calibration was also performed on the XGBoost classifier using CalibratedClassifierCV specifically using sigmoid regression to ensure that the predicted probabilities better reflected the true likelihood of an outcome and Brier score was computed to quantify the accuracy of these calibrated probabilities, with lower scores indicating better calibration.

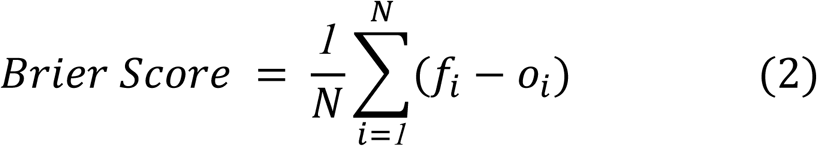

Where:

- *N* = total number of samples
- *f_i_ =* predicted probability of sample *i*
- *O_i_ =* observed outcome for sample *i* (0 = gastric cancer, 1 = non-gastric cancer)

After cross-validation, the model was refitted on the entire training dataset to optimise its performance before applying it to the test set. A Random Forest black-box classifier was trained and evaluated in the same manner as the XGBoost model. All the trained models were evaluated on the encoded test set using the trained pipeline to assess its generalisation abilities on unseen data.

### 2.11 Model Evaluation

A classification report was generated in order to provide the values of key metrics like precision, recall and F1-score. These metrics were calculated for both the positive class (gastric cancer) and the negative class (non-gastric cancer), along with their 95% confidence intervals obtained using bootstrap resampling. Specificity was explicitly reported for the negative class, while sensitivity (recall) was reported for the positive class.

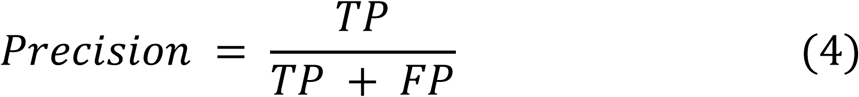

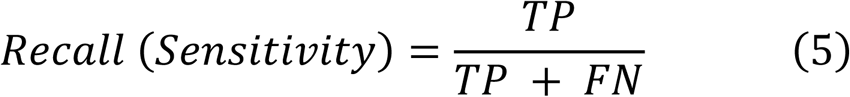

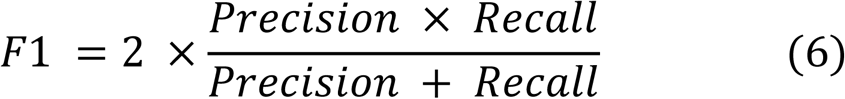

Where:

- *TP* = True Positives
- *FP* = False Positives
- *FN* = False Negatives

Along with this, the Receiver Operating Characteristic (ROC) curve was plotted to detect a trade-off between correct identification of true positives and minimization of false positives. The Area Under the Curve (AUC) score was used to summarise the ROC curve, with an AUC value closer to 1 indicating better model performance, and its 95% confidence interval was also calculated. The Area Under the Precision-Recall curve (AUPRC) was also plotted to visualise the proportion of true positives among all predicted positives against the proportion of actual positives identified by the model. The positive class was specified as gastric cancer, with the negative class being non-gastric cancer. Average precision (AP) score, computed as the area under the PR curve, offers a more effective evaluation than the ROC curve due to its sensitivity to class imbalance- the dominance of the non-gastric cancer class, and was also reported with 95% confidence interval. A higher AP score is indicative of better performance in correctly classifying the minority class. Additionally, a confusion matrix was plotted for both raw counts and percentages per cell, annotated with accuracy, sensitivity and specificity values. Calibration quality was quantified using Brier scores, with lower scores indicating that the model’s predicted probabilities were closer to the true outcome frequencies and therefore better calibrated.

### 2.12 Post-hoc Explanation

To interpret model predictions and understand the contribution of each feature, post-hoc explainability techniques were performed using SHAP Python package (version 0.47.1). The average impact of each feature on the model’s output was visualized using SHAP summary plots for the three models (Logistic Regression (baseline), XGBoost, and Random Forest). In addition to global explanations, local explanations were generated for specific test samples using SHAP waterfall plots. These plots decomposed an individual prediction into contributions from each feature relative to the model’s expected value, with the explained class chosen as either the model’s predicted class or a user-specified class index (0 for gastric cancer, 1 for non-gastric cancer). For each explained sample, the predicted class label and corresponding predicted probabilities for both classes were reported. These plots were used to illustrate the global and local feature importance across the whole dataset. These explainability techniques aid interpretation of models and help in assessing the consistency of feature contributions across different models.

## 3. Results

Classification models were developed to group subjects into the gastric cancer and non-gastric cancer classes.

Logistic Regression was used as a baseline classifier to evaluate initial predictive performance on the binary classification task of gastric cancer vs non-gastric cancer. On applying oversampling using SMOTENC, the confusion matrix (Figure 2D) revealed that the model correctly classified 61 out of 83 gastric cancer cases (73.5% sensitivity) and 161 out of 190 non-gastric cancer cases (84.7% specificity). The model produced 22 false negatives (26.5% of gastric cancer cases misclassified as non-gastric) and 29 false positives (15.3% of non-gastric cancer cases misclassified as gastric), resulting in an overall accuracy of 0.813 (95% CI: 0.766-0.864). The model displayed balanced performance across key metrics (Figure 2C), with precision values of 0.683 (95% CI: 0.591-0.784) for gastric cancer and 0.880 (95% CI: 0.831-0.928) for non-gastric cancer classification.

**Figure 2:**
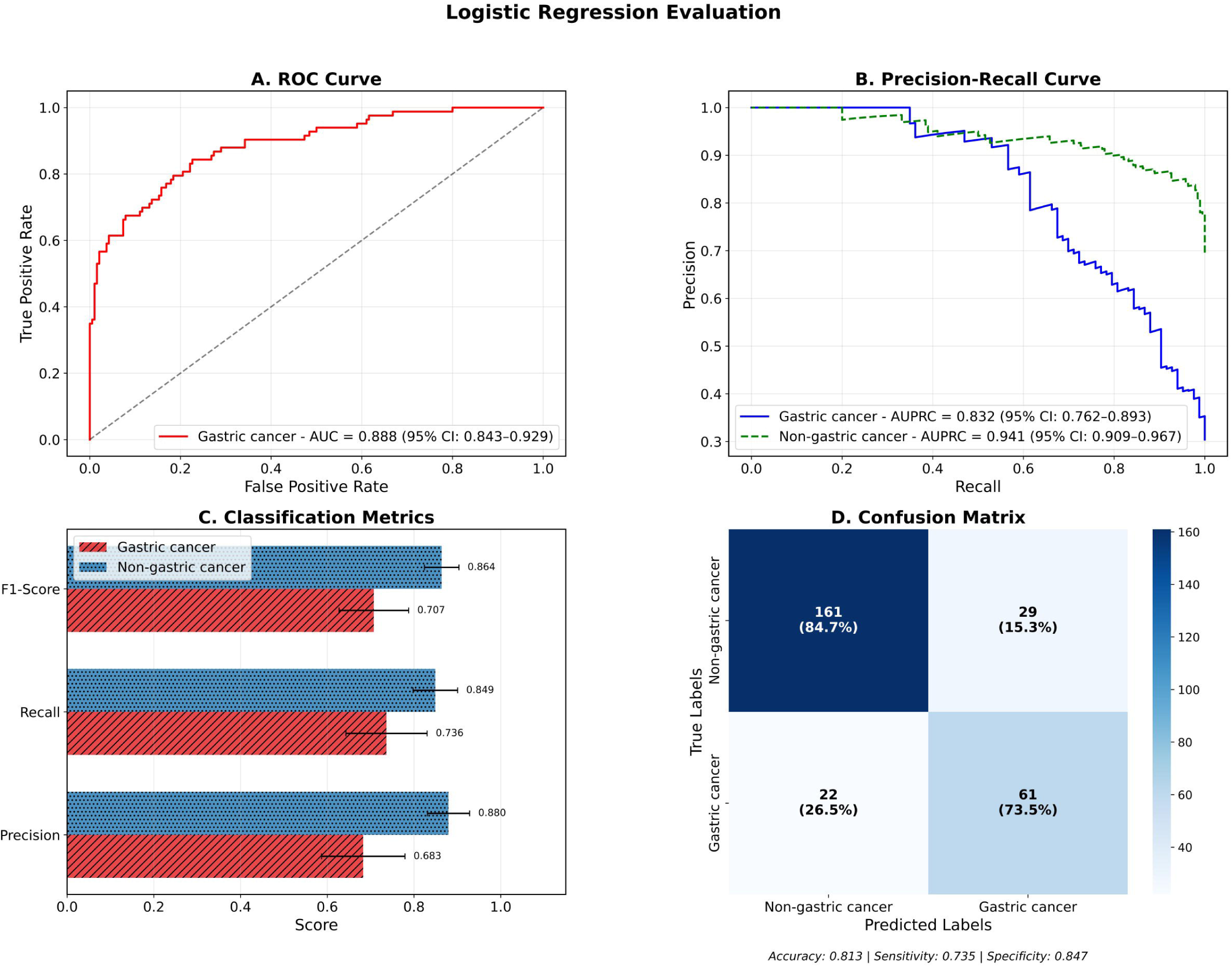
Logistic Regression model performance for gastric cancer vs non-gastric cancer classification. (A) ROC curve showing discriminative ability with AUC of 0.888 (95% CI: 0.843-0.929). (B) Precision-recall curves for both gastric cancer (AUPRC = 0.832, 95% CI: 0.762-0.893) and non-gastric cancer (AUPRC = 0.941, 95% CI: 0.909-0.967) classes. (C) Classification performance metrics comparing F1-score, recall, and precision between gastric and non-gastric cancer predictions. (D) Confusion matrix displaying true versus predicted labels with overall accuracy of 81.3%, sensitivity of 73.5%, and specificity of 84.7%.

The recall performance showed class-specific differences, with gastric cancer class recall of 0.736 (95% CI: 0.644-0.831), and non-gastric cancer recall of 0.849 (95% CI: 0.797-0.900). This resulted in F1-scores of 0.707 (95% CI: 0.623-0.784) and 0.849 (95% CI: 0.822-0.901) for gastric and non-gastric cancer classes respectively (Figure 2C). The model also achieved a good AUPRC of 0.832 (95% CI: 0.762-0.893) for gastric cancer and 0.941 (95% CI: 0.909-0.967) for non-gastric cancer classes (Figure 2B). The discriminatory power of the model denoted by an AUROC of 0.888 (95% CI: 0.843-0.929) was equal for both the classes. These findings are illustrated in Figure 2A. Detailed performance metrics with CIs are provided in Supplementary Table S6 and the classification report of the model is provided in Supplementary Table S7.

Overall, logistic regression proved to be a reasonable baseline with good performance. The lower recall for gastric cancer detection (73.6%) shows the necessity for more complex models that enhance the detection of the minority gastric-cancer class. Model calibration analysis showed reasonable probability confidence with a Brier score of 0.125 (Supplementary Figure S1).

Building on the baseline logistic regression model’s performance, we evaluated two black-box models: XGBoost and Random Forest to improve the detection of gastric cancer.

Prior to model training, feature selection was performed using ShapRFECV with cross validation to identify the most informative features for gastric cancer classification. The feature selection reduced the initial feature set from 533 to 93 features, with the optimal subset determined through 10-fold cross validation based on recall scores (Supplementary Table S8). Hyperparameter optimization was conducted using bayesian optimization with 10-fold cross validation for both the black-box models. For XGBoost, the optimal parameters were: n_estimators=1000, max_depth=7, colsample_bytree=1.0, subsample=0.601, and learning_rate=0.0279. Random Forest achieved optimal performance with n_estimators=955, max_depth=10, and min_samples_split=3. Complete hyperparameter configurations are provided in Supplementary Table S9.

XGBoost displayed superior performance over the baseline, with the confusion matrix revealing correct classification of 66 out of 83 gastric cancer cases (79.5% sensitivity) and 176 out of 190 non-gastric cancer cases (92.6% specificity). The model produced 17 false negatives (20.5% of gastric cancer cases misclassified as non-gastric) and 14 false positives (7.4% of non-gastric cancer cases misclassified as gastric), resulting in an overall accuracy of 0.886 (95% CI: 0.850-0.920) as shown in Figure 3D. The model displayed excellent performance across key metrics, with precision values of 0.829 (95% CI: 0.750-0.907) and 0.912 (95% CI: 0.871-0.950) for gastric and non-gastric cancer classes respectively. The recall performance showed improved gastric cancer detection at 0.797 (95% CI: 0.711-0.877) and maintained high non-gastric cancer recall of 0.927 (95% CI: 0.889-0.960). This resulted in F1-scores of 0.812 (95% CI: 0.745-0.871) and 0.919 (95% CI: 0.891-0.945) for gastric and non-gastric cancer classes respectively (Figure 3C). The model achieved strong AUPRC values of 0.906 (95% CI: 0.855-0.948) for gastric cancer and 0.978 (95% CI: 0.964-0.988) for non-gastric cancer classes (Figure 3B). The discriminatory power denoted by an AUROC of 0.950 (95% CI: 0.925-0.971) exceeded the baseline performance (Figure 3A).

**Figure 3:**
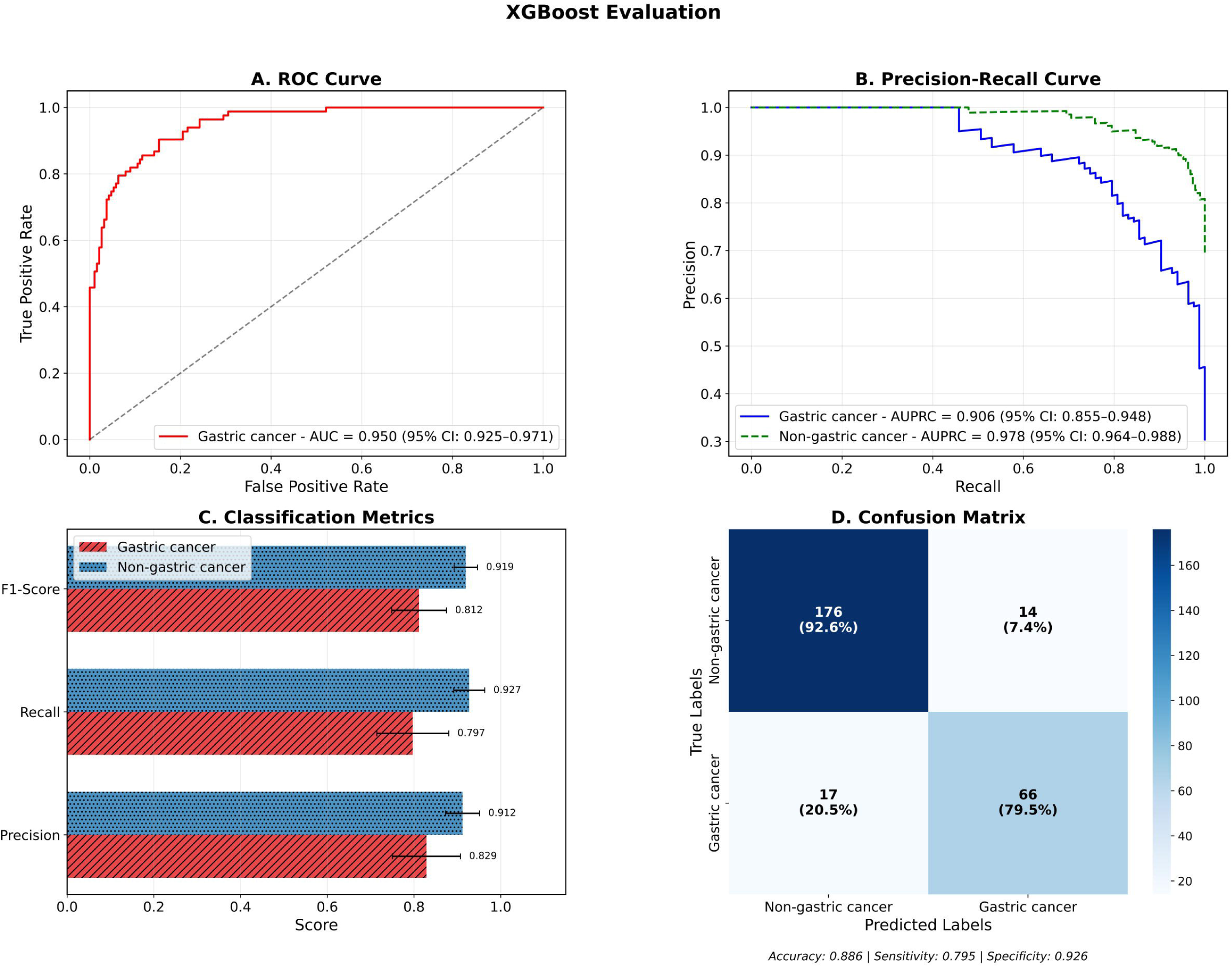
Performance evaluation of XGBoost model for gastric cancer vs non-gastric cancer classification. (A) ROC curve demonstrating superior discriminative performance with AUC of 0.950 (95% CI: 0.925-0.971). (B) Precision-recall curves showing excellent performance for both gastric cancer (AUPRC = 0.906, 95% CI: 0.855-0.948) and non-gastric cancer (AUPRC = 0.978, 95% CI: 0.964-0.988) classes. (C) Classification metrics revealing high F1-scores, recall, and precision across both classes. (D) Confusion matrix indicating improved classification accuracy of 88.6%, sensitivity of 79.5%, and specificity of 92.6%

Random Forest achieved the highest overall performance among all tested models, with the confusion matrix showing correct classification of 68 out of 83 gastric cancer cases (81.9% sensitivity) and 177 out of 190 non-gastric cancer cases (93.2% specificity). The model produced 15 false negatives (18.1% of gastric cancer cases misclassified as non-gastric) and 13 false positives (6.8% of non-gastric cancer cases misclassified as gastric), resulting in an overall accuracy of 0.897 (95% CI: 0.861-0.930) as shown in Figure 4D. The model displayed excellent balanced performance across key metrics, with precision values of 0.841 (95% CI: 0.753-0.912) for gastric cancer and 0.922 (95% CI:0.881-0.957) for non-gastric cancer classification. The recall performance displayed the highest gastric cancer detection at 0.820 (95% CI: 0.734-0.896) and robust non-gastric cancer recall of 0.932 (95% CI: 0.897-0.963). This resulted in F1-scores of 0.830 (95% CI: 0.767-0.886) and 0.926 (95% CI: 0.898-0.951) for gastric and non-gastric cancer classes respectively (Figure 4C). The model achieved excellent AUPRC values of 0.926 (95% CI: 0.888-0.959) for gastric cancer and 0.978 (95% CI: 0.961-0.990) for non-gastric cancer classes (Figure 4B). The discriminatory power denoted by an AUROC of 0.954 (95% CI: 0.929-0.976) was the highest among all tested models (Figure 4A). Detailed performance metrics with confidence intervals are provided in Supplementary Table S6 and the classification report is provided in Supplementary Table S7.

**Figure 4:**
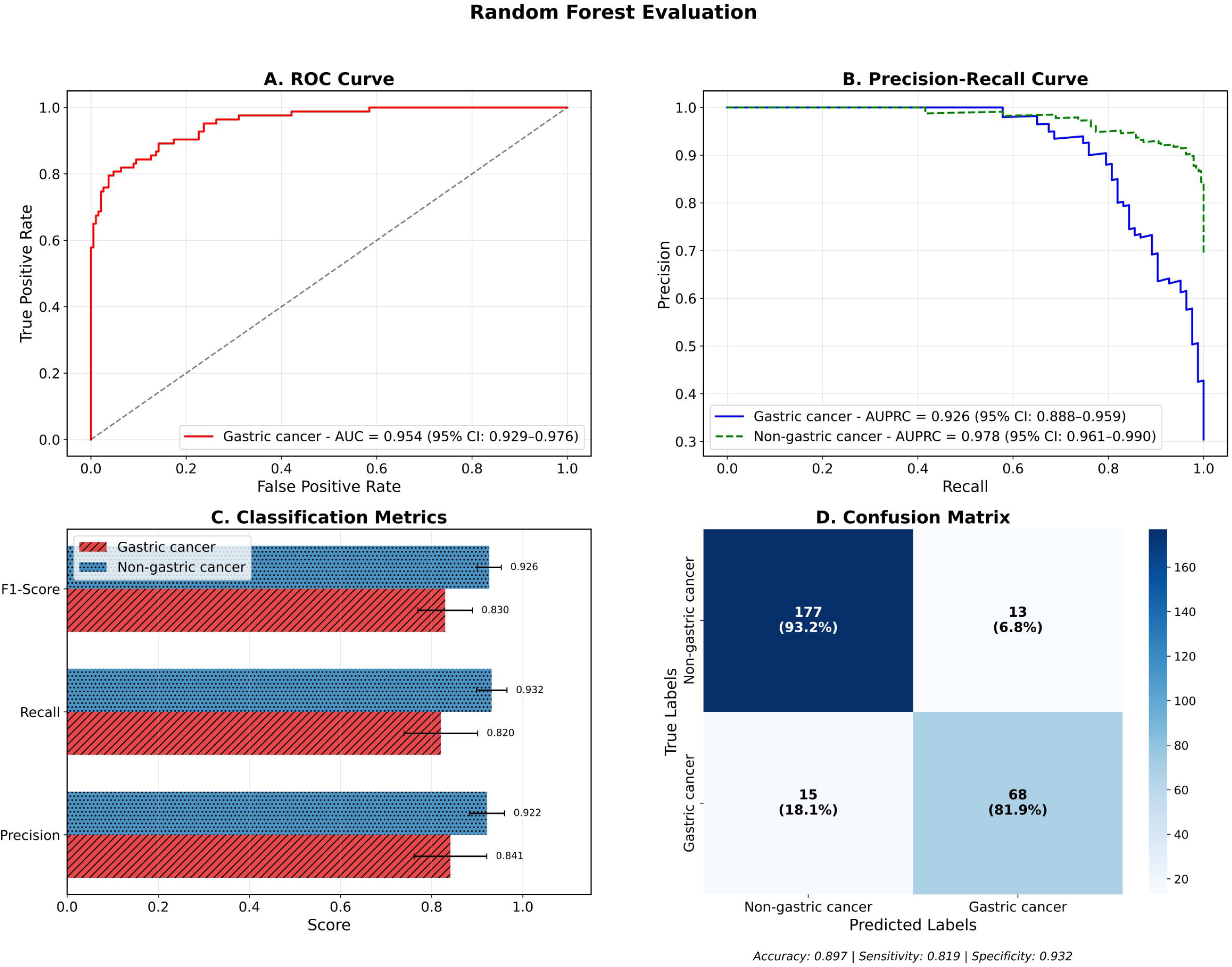
Performance evaluation of Random Forest model for gastric cancer vs non-gastric cancer classification. (A) ROC curve showing excellent discriminative capacity with AUC of 0.954 (95% CI: 0.929-0.976). (B) Precision-recall curves demonstrating robust performance for gastric cancer (AUPRC = 0.926, 95% CI: 0.888-0.959) and non-gastric cancer (AUPRC = 0.978, 95% CI: 0.961-0.990) classification. (C) Classification performance metrics indicating balanced precision and recall across both classes. (D) Confusion matrix showing the highest overall accuracy of 89.7%, with sensitivity of 81.9% and specificity of 93.2%.

Both black-box models outperformed the baseline, with Random Forest achieving superior performance in gastric cancer detection (sensitivity: 82.0% vs 79.7% for XGBoost vs 73.6% for baseline) and overall accuracy (89.7% vs 88.6% vs 81.3% respectively). Model calibration analysis revealed improved probability confidence for both black-box models, with XGBoost achieving a Brier score of 0.087 and Random Forest achieving 0.078 after calibration, representing 30% and 38% improvements over the baseline score of 0.125 respectively (Supplementary Figure S2 and S3). The superior performance of both black-box models, particularly Random Forest’s enhanced gastric cancer detection capability, demonstrates the effectiveness of tree-based algorithms for this complex classification task.

DeLong’s test was performed to statistically evaluate the differences between the areas under two ROC curves derived from distinct models. The difference in the AUROC of XGBoostClassifier and the baseline Logistic Regression was statistically significant with a p-value equal to 0.0017 and Z score equal to −3.13. Similarly, the difference in AUROC of Random Forest and the baseline Logistic Regression was statistically significant with a p-value of 0.0013 and Z score of −3.20. Both black-box models displayed statistically significant improvements over the baseline logistic regression model.

### 3.1 Model Explanations

The global SHAP summary plots provide post-hoc global explanations of the model to identify the features that contributed most to the model’s predictions.

In the case of the baseline logistic regression model, the model primarily relied on demographic variables, with Age as the most influential predictor, followed by Continent (Asia and Europe). Aggregate variant categories such as start_retained_variant, intergenic_region and other combinations of different variant categories also contributed, though with moderate impact (Figure 5A). Sequence-derived features such as k-mers, Z-curve were not in the top 10 influential features for the baseline model.

**Figure 5.**
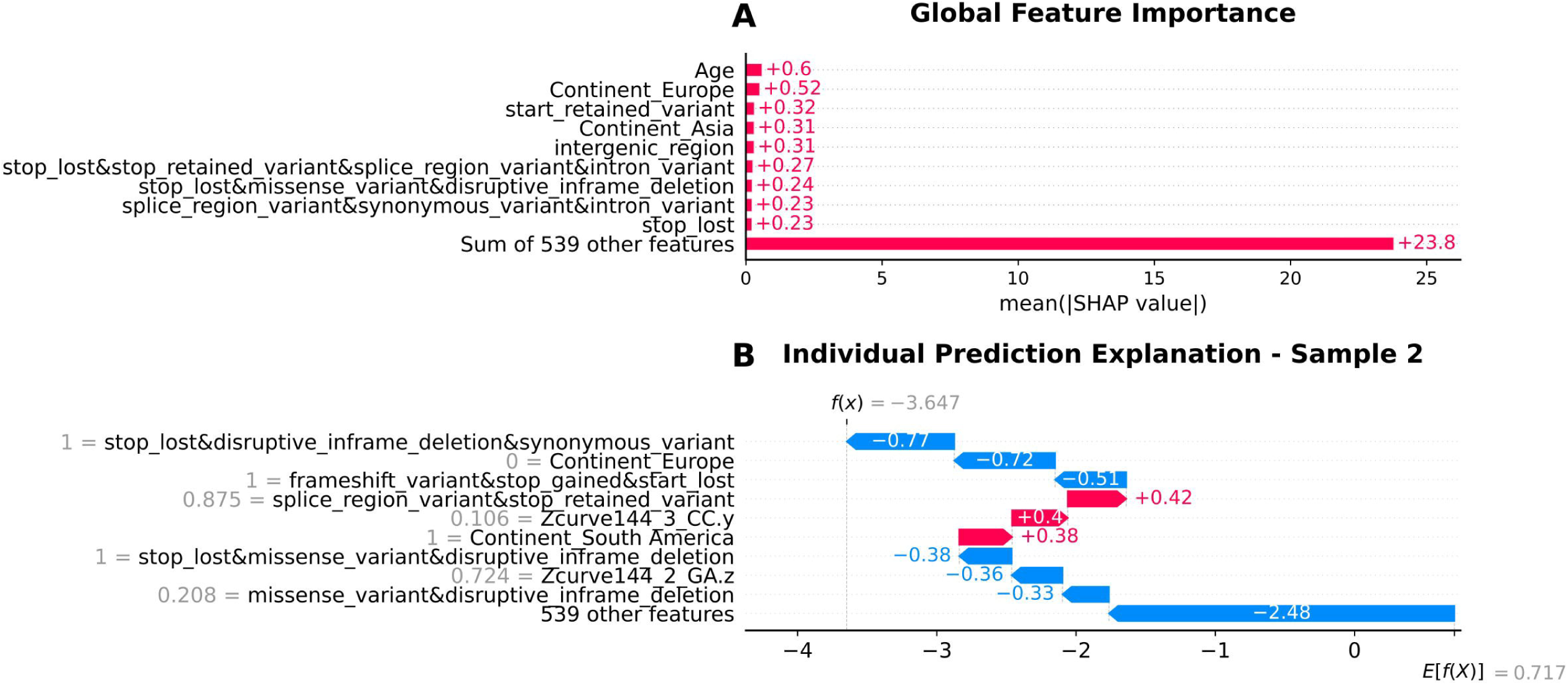
SHAP (SHapley Additive exPlanations) analysis of logistic regression model interpretability. (A) Global feature importance showing the mean absolute SHAP values across all features, with Age (0.60) and Continent_Europe (0.52) as the most influential predictors, while 539 additional features collectively contribute 23.8 to model predictions. (B) Individual prediction explanation for Sample 2 (f(x) = −3.647, E[f(X)] = 0.717) demonstrating how specific feature values contribute toward non-gastric cancer prediction (red bars, positive values) or gastric cancer prediction (blue bars, negative values), with the overall negative prediction score indicating classification toward gastric cancer (class 0).

The XGBoost model captured a combination of demographic, aggregate variant categories and sequence-derived features. Age was the top predictor here too followed by a sequence-derived feature CAC k-mer. Among variant categories, frameshift_variant and frameshift_variant&stop_lost had considerable contributions, while other sequence k-mers such as AGC, ATA, and GTA also ranked highly. In addition to the above mentioned features, Z-curve descriptors (Zcurve144_1_CA.y and Zcurve144_3_AA.z) appeared among the top 10 influential predictors, suggesting that genome composition features provided additional discriminatory power (Figure 6A).

**Figure 6.**
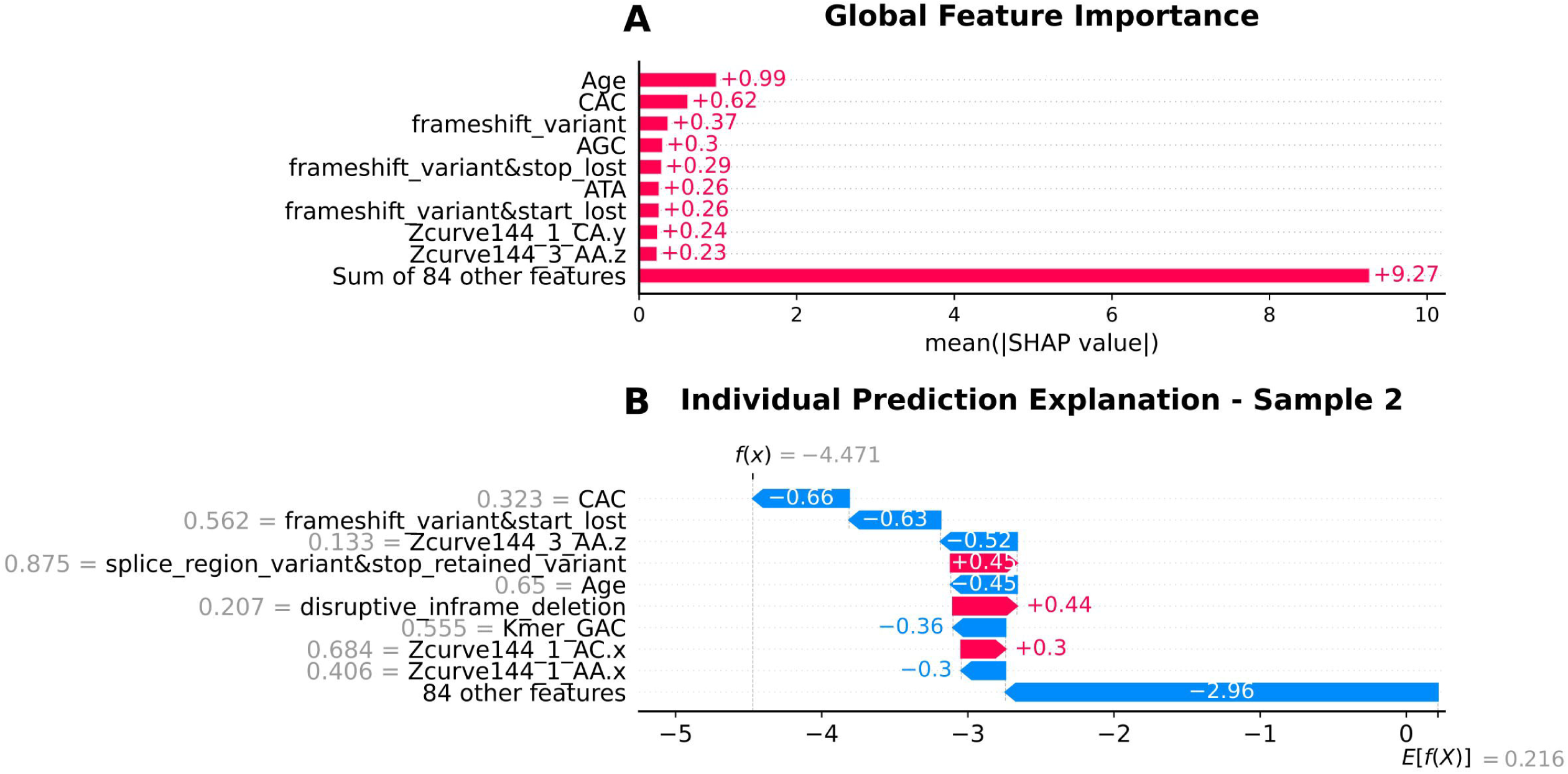
SHAP analysis revealing feature importance and prediction explanations for the XGBoost model. (A) Global feature importance highlighting Age (0.99) as the dominant predictor, followed by CAC (0.62) and frameshift_variant (0.37), with 84 other features contributing 9.27 collectively. (B) Individual prediction explanation for Sample 2 (f(x) = −4.471, E[f(X)] = 0.216) showing how CAC (−0.66), frameshift variants (−0.63), and age-related features (−0.52) drive the prediction strongly toward gastric cancer (class 0), while some variants categories provide modest contributions toward non-gastric cancer classification.

Similar to XGBoost, Age and CAC k-mer were the top predictors, indicating their consistent importance across models. The Random Forest also emphasized sequence descriptors, such as MMI_AT, CAA, and CKSNAP_TA.gap2, alongside variant categories like frameshift_variant&stop_lost&splice_region_variantstop_retained_variant and disruptive_inframe_deletion&synonymous_variant. Additional k-mers (ATA, AGA) and structural variant categories (for e.g., conservative_inframe_insertion) contributed, though with smaller SHAP values compared to Age and CAC (Figure 7A).

**Figure 7.**
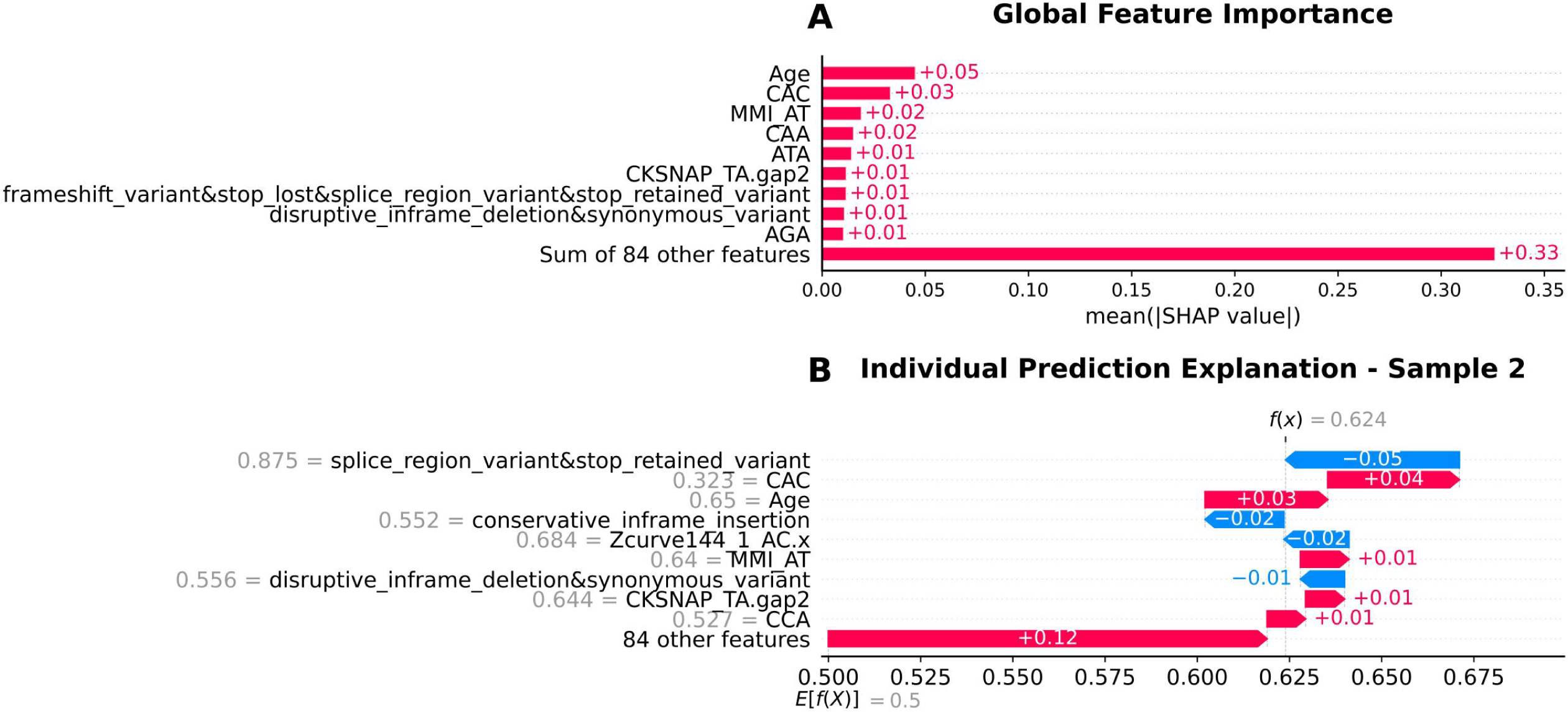
SHAP analysis demonstrating the interpretability of the Random Forest model predictions. (A) Global feature importance showing relatively balanced feature contributions with Age (0.05), CAC (0.03), and MMI_AT (0.02) as leading predictors, and 84 other features contributing 0.33 collectively, indicating more distributed feature utilization compared to other models. (B) Individual prediction explanation for Sample 2 (f(x) = 0.624, E[f(X)] = 0.5) revealing subtle feature contributions, with splice_region_variant providing influence toward gastric cancer (−0.05) and CAC contributing toward non-gastric cancer (+0.04), with the overall positive prediction score indicating classification toward non-gastric cancer (class 1).

Across all models, age consistently emerged as the most influential predictor, while Continent_Europe was particularly important in the Logistic Regression model. Tree-based classifiers (XGBoost and Random Forest) captured a broader range of sequence k-mers and variant categories, suggesting they were better able to integrate complex genomic patterns along with demographic features.

To interpret individual-level predictions, we generated SHAP waterfall plots for representative test samples that were classified as gastric cancer (positive class, encoded as 0) by each model. These plots illustrate how genomic and demographic features contributed positively or negatively to the probability of gastric cancer in correctly predicted cases.

The logistic regression model relied heavily on combined variant types, with stop_lost&disruptive_inframe_deletion&synonymous_variant and geographic factor (patient not being from Continent_Europe) as primary features that contribute to gastric cancer classification. The 539 aggregated features contributed substantially, while splice_region_variant&stop_retained_variant provided the strongest opposing influence (Figure 5B).

The XGBoost model emphasized sequence composition, with CAC and frameshift_variant&start_lost as top contributors toward gastric cancer prediction. Z-curve parameter, Zcurve144_3_AA.z, age, and k-mer GAC played a significant role. The 84 aggregated features showed strong gastric cancer contribution, while splice_region_variant&stop_retained_variant, disruptive_inframe_deletion, and Zcurve144_1_AC.x contributed to the non-gastric cancer class (Figure 6B).

The Random Forest model showed variant categories pushing the classification towards the gastric cancer class with splice_region_variant&stop_retained_variant being a comparatively strong contributor, while the remaining variant categories such as conservative_inframe_insertion and disruptive_inframe_deletion&synonymous_variant show moderate contributions. Genome composition features like Z-curve descriptors (Zcurve144_1_AC.x) also contributed to the gastric cancer class (Figure 7B).

## 4. Discussion

Three machine learning algorithms were used to identify primarily genomic variables that contributed to the development of gastric cancer in individuals infected with *H. pylori*. In order to be inclusive and accessible to clinicians, white-box models were used in addition to high-performance black-box machine learning models. A variety of explainability measures like SHAP were also utilized to explain identified associations and increase clinician confidence, and this framework was designed to uncover dynamic genomic risk factors and supplement clinician decision-making. This is demonstrably superior to traditional modeling methods which are constrained by the number of variables that can be incorporated into the study and unable to unearth complex associations.

It can be seen that age was consistently identified as the top predictor of gastric cancer across all three models. This aligns with existing scientific literature implicating older patients as more susceptible to developing gastric cancer, with around 6 out of every 10 people diagnosed with gastric cancer being 65 years or older^30^.

The top 3 predictors in the baseline model were age, belonging to Europe and silent mutations in the start codons (Figure 5A). Fatality linked to gastric cancer has been shown to be particularly high in regions of East Asia and Eastern Europe^31^. The malignancy of certain geographical strains could be in part due to differences in virulence genotypes like *cagA* motifs^32^. The silent mutations in the start codon could lead to the production of proteins with altered functional properties, thus influencing the malignancy of the *H. pylori* strain.

The top 3 contributing factors in the XGBoost model were age, the reverse kmer CAC and the simultaneous presence of frameshift mutations and loss of stop codons (Figure 6A). The presence of the trinucleotide CAC could be indicative of distinctive pathogenicity islands or similar genomic signatures associated with *H. pylori* infection and adaptation. Stop codons have also been implicated in existing literature, with a study depicting the loss of expression of the virulence gene BabA due to a mutation resulting in a stop codon being acquired^33^. The loss of stop codons could be likely to capture phase variation events that control the expression of key infection factors and thus influencing gastric cancer risk.

The top 3 contributing factors in the Random Forest model were age, the reverse kmer CAC and the presence of AT rich regions (Figure 7A). While the first two features were also the most important in the XGBoost model, multivariate mutual information aids in capturing higher-order nucleotide correlations beyond regular k-mer counts. The high correlation with AT could indicate the difference in the degree of sequence-order through structured repeats and conserved motifs between cancer-associated and benign strains.

The two black-box models also consistently identified the presence of the CAC, ATA and AAA reverse kmers as associated with increased discriminatory power between infection outcomes. Combinations of variants have also been marked as gastric-cancer inducing factors in both the XGBoost and Random Forest models, such as the high presence of all 4 mutations-frameshift variants, loss of stop codons, missense variants and splice region variants. These variants result in an alteration of amino acids in bacterial proteins, and could reflect divergent strains, altered virulence factors and genetic plasticity. Previous studies have found that many common allelic variants are more frequent in outer-membrane associated virulence genes, like *sabA* and *babA*^34^. Besides corroborating existing literature on the causative risk factors of malignant *H. pylori* infection induced gastric cancer and thus establishing itself as reliable, our study can also advance research that seeks to detect and mitigate *H. pylori* outcomes through personalized treatment approaches. Further studies that seek to investigate causal inference between identified features can also make use of our explainable machine learning framework. Although the models possessed remarkable discriminatory power, a variety of pertinent features were not incorporated into the model. It has been shown that family history, smoking habits, diet and other clinical features play a very important role in the development of *H. pylori* infection associated outcomes, along with the role of host immune factors^35^. The data also had a large number of European samples (870), and since the data was publicly available, there could have been varying biases introduced at different stages of data collection like participation bias, and the lack of inclusion of control patients without *H. pylori* infection. Access to more diverse data sources, particularly from the Global South, could mitigate existing geographic bias and render the model more generalizable.

## 5. Conclusions

Integrating machine learning with bedside treatment will be a significant stride in early detection and personalized diagnostic methods for identifying the severity of *H. pylori* infection responses. In an attempt to improve clinical trust in black-box models, explainable AI was used to break down top features to improve transparency. Among the evaluated models, the two black-box models achieved greater recall in the prediction of gastric cancer compared to the baseline logistic regression model. The top predictor was consistently found to be patient age across all three models, while different genomic markers were identified by each of the models.

Incorporating genomic features of the host along with clinical metadata could enhance the predictive power and the robustness of the models since current *H. pylori* prediction models focus largely on clinical features. As gene expression plays a major role in the development of infection, integration of transcriptomic data and host gut metagenomic data could account for gene expression and interaction of *H. pylori* with other gut microbes. The sequence-derived genomic features implicated in this study open avenues for further research into the *H. pylori* genome beyond the already implicated virulence genes to assess possible causation behind the associations found in this study. Assimilating real-time data gathering options for the model through collaboration with medical centres, incorporating clinical feedback loops and refining the user interface can render it a powerful tool for spontaneous risk-assessment and bolster regional health infrastructure. The high accuracy of our research model in discriminating between severe and benign *H. pylori* infection outcomes underscore its potential impact in deploying it in resource-limited populations that are disproportionately plagued by *H. pylori* infections.

## Declarations

### Ethics approval and consent to participate

Not applicable

### Consent for publication

Not applicable

### Availability of data and materials

The *Helicobacter pylori* genomes analyzed in this study are publicly available on the National Center for Biotechnology Information (NCBI) GenBank repository (https://www.ncbi.nlm.nih.gov/genbank/) and EnteroBase (https://enterobase.warwick.ac.uk/). Accession numbers for all 1,363 genomes used in analyses are provided in Supplementary Table 1. The custom code and analysis pipelines used in this study are available on GitHub (https://github.com/Venkatesh-99/HP_ML/) and archived in Zenodo at (https://doi.org/10.5281/zenodo.17085927).

### Author Contributions Statement

SPW and VN contributed equally to conceptualization, data curation, investigation and methodology design, software and formal analysis, data visualization, validation, writing the original draft, reviewing and editing the manuscript. MPT contributed to data curation and formal analysis. JJJ, NV and GGT contributed to conceptualization, reviewing and editing the manuscript. BV contributed to reviewing and editing the manuscript, provision of resources, project administration and supervision. All authors read and approved the final manuscript.

## Acknowledgements

Not applicable

## Additional files

**File name:**

Additional file 1

**File format:**

.xlsx (Microsoft Excel Spreadsheet)

**Title of data:**

Supplementary Tables S1-S9: Features used, detailed model metrics, optimal hyperparameters

**Description of data:**

- Supplementary Table S1: Clinical metadata of the patients. Includes patient demographics like their age, sex, their geographical location and the disease phenotype.
- Supplementary Table S2: Presence/absence data of *H. pylori* virulence genes implicated in gastric cancer
- Supplementary Table S3: Sequence-derived features extracted from *H. pylori* genomes using iFeatureOmega and MathFeature
- Supplementary Table S4: Aggregate variant categories of *H. pylori* genomes
- Supplementary Table S5: The final all feature-types integrated dataset
- Supplementary Table S6: Detailed evaluation metrics for all three models with class-specific precision, recall, F1-score, AUROC and AUPRC scores with lower and upper confidence intervals
- Supplementary Table S7: Classification reports of all three models
- Supplementary Table S8: ShapRFECV feature elimination report. Contains information about the selected and eliminated features based on the recall metric
- Supplementary Table S9: Optimal hyperparameters selected using Bayesian optimization for XGBoost and Random Forest models

**File name:**

Additional file 2

**File format:**

.png (Portable Network Graphics image)

**Title of data:**

Calibration plot of logistic regression classifier

**Description of data:**

This figure shows the calibration performance of the logistic regression model for predicting gastric cancer. The fraction of positives is plotted against the mean predicted probability, with the dashed diagonal line representing perfect calibration. The Brier score is provided as a summary measure of calibration accuracy.

**File name**

Additional file 3

**File format**

.png (Portable Network Graphics image)

**Title of data**

Calibration plots of extreme gradient boosting (XGBoost) classifier before and after calibration

**Description of data**

This figure presents side-by-side calibration plots of the XGBoost classifier before and after calibration. Each plot shows the fraction of positives against mean predicted probability, with the dashed diagonal line representing perfect calibration. Brier scores before and after calibration are included to quantify improvement in model calibration.

**File name**

Additional file 4

**File format**

.png (Portable Network Graphics image)

**Title of data**

Calibration plots of random forest classifier before and after calibration

**Description of data**

This figure shows the calibration performance of the random forest classifier, with separate panels for before and after calibration. Each plot compares predicted probabilities with observed outcomes, with the diagonal line indicating perfect calibration. Brier scores are reported to demonstrate changes in calibration accuracy following adjustme

## 6. List of abbreviations

*H. pylori*: *Helicobacter pylori*
XGBoost: eXtreme Gradient Boosting
SMOTE-NC: Synthetic Minority Over-sampling Technique for Nominal and Continuous
SHAP: SHapley Additive exPlanations
AUROC: Area Under Receiver Operating Characteristic curve
MALT: Mucosa-associated lymphoid tissue
WHO: World Health Organisation
*cagA*: cytotoxin-associated gene A
cagPAI: cag pathogenicity island
*vacA*: vacuolating cytotoxin A
BabA: blood group antigen binding adhesin
*oipA*: outer inflammatory protein A
sLe^X^: sialylated Lewis antigens
SRA: Sequence Read Archive
NAC: Nucleic Acid Composition
MMI: Multivariate Mutual Information
ROC: Receiver Operating Characteristic
AUC: Area Under the Curve
AUPRC: Area Under the Precision-Recall curve
AP: Average precision

