## Supplementary figures and images for "Predicting Clinical Outcomes in *Helicobacter pylori-*positive Patients using Supervised Learning through the Integration of Demographic and Genomic Features"

### Supplementary Figure 1

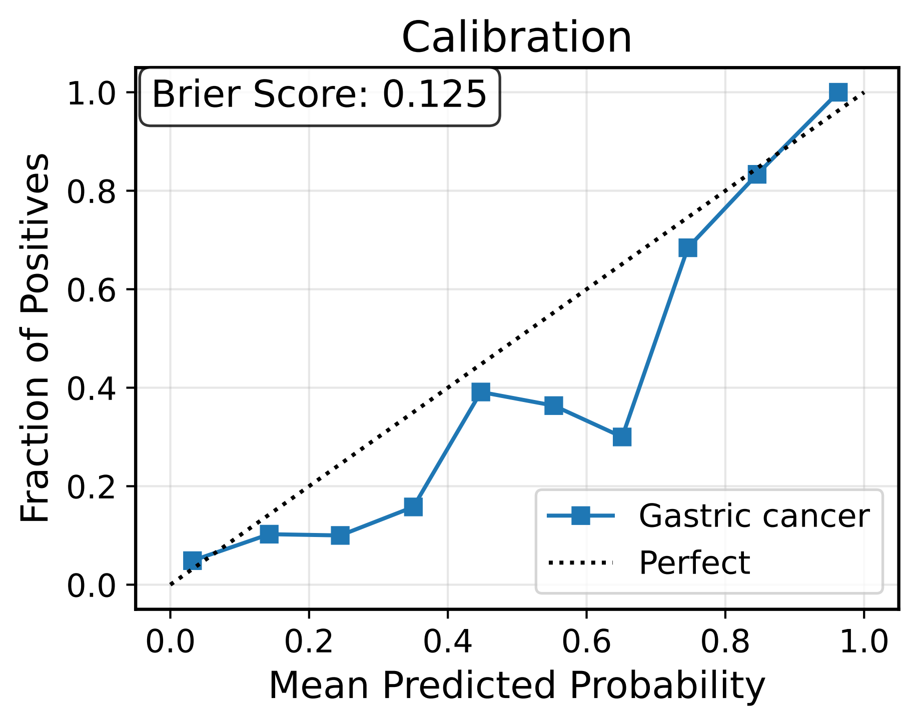

### Supplementary Figure 2

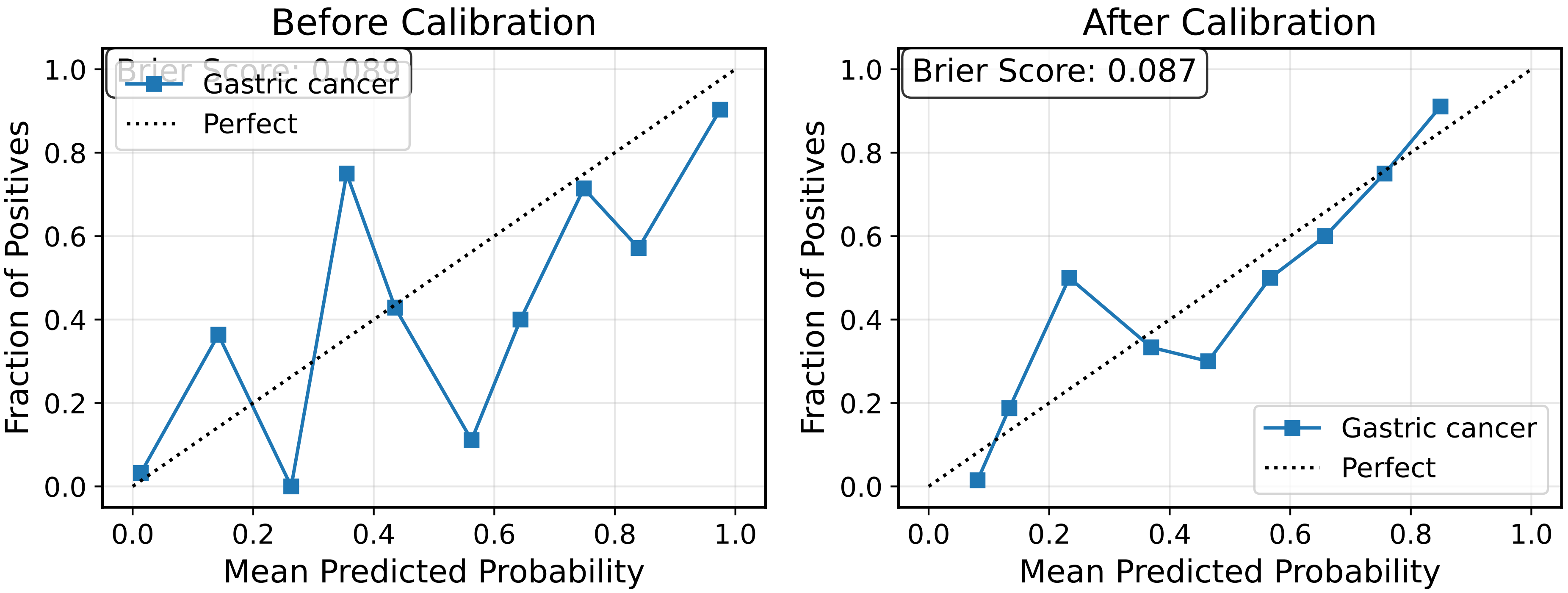

### Supplementary Figure 3

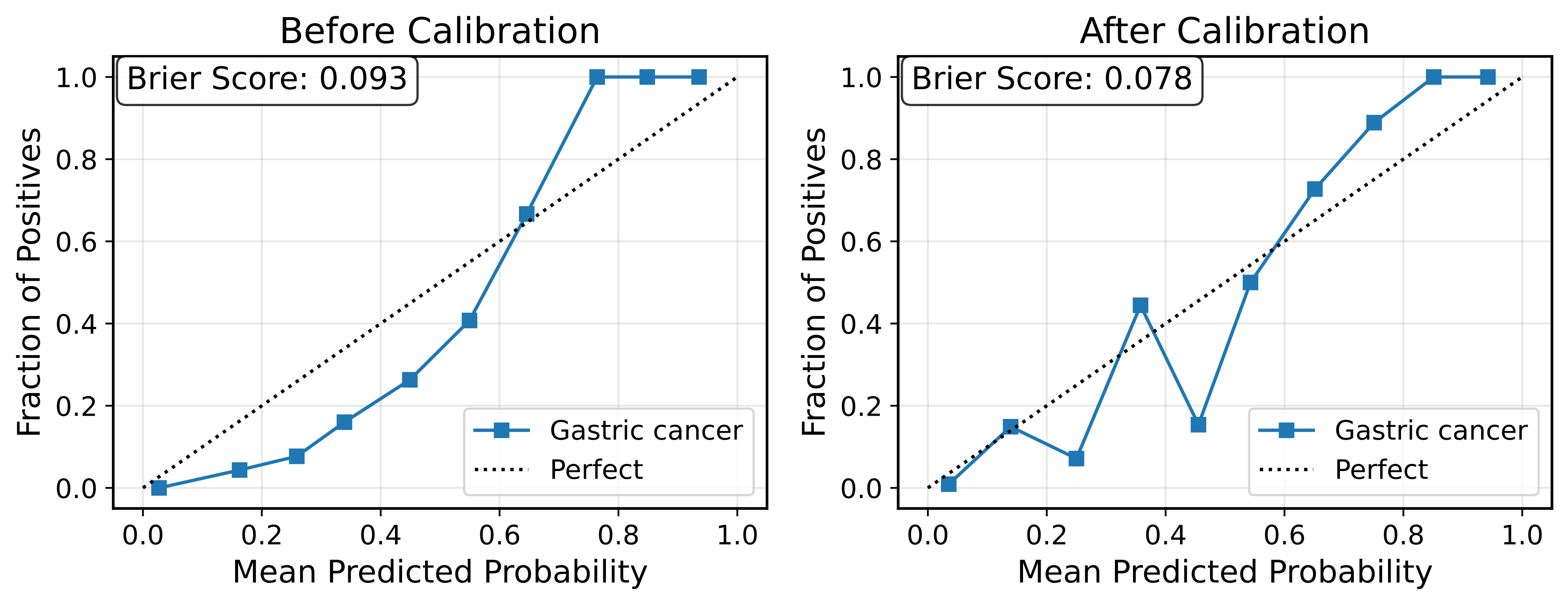
